# Chronic alcohol consumption enhances the differentiation capacity of hematopoietic stem and progenitor cells into osteoclast precursors

**DOI:** 10.1101/2025.02.05.636743

**Authors:** Hami Hemati, Madison B. Blanton, Jude koura, Rupak Khadka, Kathleen A. Grant, Ilhem Messaoudi

**Affiliations:** Microbiology, Immunology and Molecular Genetics, College of Medicine, University of Kentucky, Lexington, KY, United States; Pharmaceutical Sciences, College of Pharmacy, University of Kentucky, Lexington, KY, United States; Division of Neuroscience, Oregon National Primate Research Center, Oregon Health and Science University, OR, United States

**Keywords:** Chronic alcohol consumption, Hematopoietic stem and progenitor cells, Osteoclastogenesis, Osteoclast precursors, Osteoclast, Osteoporosis

## Abstract

Chronic alcohol consumption (CAC) is associated with an enhanced risk of bone fracture, reduced bone density, and osteoporosis. We have previously shown using a rhesus macaque model of voluntary ethanol consumption that CAC induces functional, transcriptomic, and epigenomic changes in hematopoietic stem and progenitor cells (HSPCs) and their resultant monocytes/macrophages, skewing them towards a hyper-inflammatory response. Here, we extended those studies and investigated alterations in osteoclasts, which, in postnatal life, are differentiated from HSPCs and play a critical role in maintaining bone homeostasis. Analysis using spectral flow cytometry revealed a skewing of HSPCs towards granulocyte-monocyte progenitors (GMPs) with the CAC group that was in concordance with an increased number of colony-forming unit-granulocyte/macrophage (CFU-GM). Additionally, HSPCs from animals in the CAC group incubated with M-CSF and RANKL were more likely to differentiate into osteoclasts, as evidenced by increased Tartrate-Resistant Acid Phosphatase (TRAP) staining and bone resorption activity. Moreover, single-cell RNA sequencing of differentiated HSPCs identified three clusters of osteoclast precursors in the CAC group with enhanced gene expression in pathways associated with cellular response to stimuli, membrane trafficking, and vesicle-mediated transport. Collectively, these data show that CAC-derived hematopoietic progenitor cells exhibit a higher capacity to differentiate into osteoclast precursors. These findings provide critical insights for future research on the mechanisms by which CAC disrupts monopoiesis homeostasis and enhances osteoclast precursors, thereby contributing to reduced bone density.

## INTRODUCTION

Alcohol misuse is prevalent in the United States, with 29.5 million individuals over the age of 12 suffering from alcohol use disorder (AUD) in 2022^1^. AUD is associated with numerous detrimental health effects, such as a higher occurrence and severity of fractures, reduced bone density, and osteoporosis^2–5^. A series of meta-analyses have demonstrated that in contrast to light/moderate alcohol consumption^2^, chronic heavy alcohol consumption is associated with a reduction in bone mineral density (BMD) and an increased risk of bone fractures^2, 3, 5^. Specifically, there is a higher risk of hip fractures in individuals consuming three or more standard drinks of alcohol per day^3^. Furthermore, individuals who consume two or more alcoholic beverages per day are at increased risk of developing osteoporosis compared to abstainers^5^. Additionally, prenatal alcohol exposure may contribute to lowering height and BMD in early adulthood^6^. Given the significant impact of alcohol on bone, it is imperative to investigate the underlying mechanisms by which alcohol affects generation and the activity of cells integral to bone health.

Normal bone remodeling relies on the dynamic equilibrium between osteoclastic resorption and osteoblastic formation. In postnatal life, osteoclasts originate from hematopoietic stem cell-derived monocyte/macrophage precursor cells^7, 8^. Differentiation of HSPCs to osteoclasts relies on RANK/RANKL and M-CSF/CSF1R signaling^8^. This results in the formation of multinucleated mature osteoclasts with varying nuclei numbers and cell sizes^9^. Diminished osteoblastic bone formation or excessive osteoclastogenesis was reported in various pathological conditions, including osteoporosis, autoimmune arthritis^10^, periodontitis^11^, and bone metastasis^12^. The underlying common thread in these pathological conditions is an inflammatory process that leads to the accelerated differentiation of osteoclast precursors into mature osteoclasts^13–15^.

Monocyte lineage is a potential osteoclast precursor^7, 8^. Our previous research has demonstrated that CAC skews the differentiation of HSPCs toward monocyte and granulocyte progenitors^16^. These observations were consistent with extensive epigenetic modifications associated with increased chromatin accessibility at promoters and intergenic regions that regulate inflammatory responses observed in monocytes and their progenitors^16^. Therefore, the impact of CAC on HSPCs may extend to osteoclasts. However, the molecular mechanisms by which CAC alters HSPCs and, subsequently, osteoclast precursors remain poorly understood. In this study, we used a rhesus macaque model of voluntary ethanol self-administration that accurately resembles human physiology in response to alcohol consumption. Alcohol consumption in this model was reported to be associated with reduced bone formation^17^, suppression of bone remodeling^18^, reduced intracortical bone porosity^19^, and decreased bone turnover^20, 21^. To further explore the impact of CAC on bone health at the cellular level, we investigated the impact of CAC on HSPCs and their differentiation into osteoclast precursors using phenotypic and functional readouts and scRNA-seq. Furthermore, we explored the potential impact of CAC-induced osteoclastogenesis on the differentiation of HSPCs toward monocyte and granulocyte progenitors. Collectively, our findings contribute to a deeper understanding of how alcohol consumption adversely affects bone health through the alteration in differentiation dynamics of hematopoietic stem cells and progenitors.

## MATERIALS AND METHODS

### Animals and bone marrow samples

This study used samples obtained through the Monkey Alcohol Tissue Research Resource (www.matrr.com) from a rhesus macaque model of voluntary ethanol consumption^22–24^. Briefly, the experimental design involved a 90-day induction period during which the subjects were acclimated to a 4% w/v ethanol solution. Following this phase, the animals were given the choice to consume either a 4% w/v ethanol solution or water for 22 hours per day for one year. In this model, animals naturally divide into heavy and moderate drinkers within 2 to 3 months, and these drinking patterns remain consistent for up to 12 months. This protocol results in approximately 40% of the animals transitioning into heavy or very heavy drinkers^25, 26^. Bone marrow samples were obtained from control (n=8: 4 females and 4 males, from cohorts 7b, 6a, and 9) and macaques that engaged in very heavy drinking throughout the 12-month ethanol exposure (n=9: 5 females and 4 males from cohorts 7a, 6a, and 17). Ethical approval for this study was granted by the Institutional Animal Care and Use Committee of the Oregon National Primate Research Center (ONPRC).

### *In vitro* osteoclastogenesis assay and TRAP staining

Cryopreserved bone marrow samples were thawed in a 37°C water bath, followed by washing them with RPMI 1640 (Gibco) supplemented with 10% fetal bovine serum (FBS), 1% L-glutamine (GeminiBio), 1% penicillin-streptomycin (GeminiBio), and 1% DNase 1 (Thermofisher Scientific) and centrifugation at 300×g for 5 minutes. Cells were then resuspended in RPMI 1640 with 10% FBS, 1% L-glutamine, and 1% penicillin-streptomycin and counted. The bone marrow mononuclear cells were incubated in MEMα with GlutaMAX™ Supplement (Gibco), supplemented with 10% FBS, 1% penicillin-streptomycin, and human M-CSF (10 ng/ml final concentration; Peprotech) overnight at 37°C in a 5% CO₂ atmosphere. Non-adherent cells were collected by gently removing the supernatant and centrifuging at 300×g for 3 minutes. Cells were then plated at a density of 0.4×10⁶ cells/well in 48-well plates in induction media consisting of MEMα supplemented with 10% FBS, 1% penicillin-streptomycin, 30 ng/mL M-CSF, and 80 ng/mL RANKL. Induction media was replaced every 48 hours. 24 hours after the third media change, the culture supernatant was discarded, and cells were fixed for 5 minutes. Tartrate-resistant acid phosphatase (TRAP) staining was performed using a TRAP kit per manufacturer recommendations (Takarabio). Briefly, 200 µL aliquot of the TRAP substrate solution was added to each well, and plates were incubated at 37°C for 30 minutes. The substrate solution was then removed, and wells were washed three times with sterile distilled water. The whole well area was imaged to detect TRAP activity as purplish-red staining in osteoclasts. Images were analyzed using ImageJ software. Osteoclasts were defined as TRAP-positive cells ≥3 nuclei.

### Bone resorption assay

The bone resorption activity was performed on a fluoresceinated calcium phosphate-coated plate using a bone resorption assay kit (Cosmo Bio) following manufacturer instructions. Briefly, 0.1 ml of fluoresceinamine-labeled chondroitin sulfate (FACS) was added to the calcium phosphate-coated plates under aseptic conditions, ensuring minimal disturbance to the coating. Plates were incubated at 37°C for 2 hours in the dark, then washed twice with 0.2 ml of PBS. Non-adherent bone marrow cells collected as described above were seeded onto the FACS-coated plates at a density of 0.2 × 10^6^ cells/well in a phenol red-free MEMα supplemented with 10% FBS, 1% penicillin-streptomycin, 1% GlutaMax (ThermoFisher), 30 ng/ml human M-CSF, and 100 ng/ml human RANKL. Following the kit’s recommendation and our optimization assays, we used a higher concentration of RANKL than when culturing cells on plastic plates in the osteoclastogenesis assay. The plates were maintained in light-shielded conditions throughout the experiment. The condition media was collected and replaced every 48 hours. The fluorescence intensity of the collected conditioned media was measured using a plate reader at an excitation wavelength of 485 nm and an emission wavelength of 535 nm. Twenty-four hours after the third media change, the wells were treated with 0.2 ml of 5% sodium hypochlorite for 5 minutes to remove adherent cells. Plates were then washed with water, dried, and stored for subsequent imaging. The pits, representing resorption activity, were analyzed using ImageJ software. Images were converted to 8-bit grayscale and were thresholded with “Dark background” selected. The threshold was automatically applied to highlight pits.

### Single RNA-seq library preparation and data analysis

The cells were collected following osteoclastogenesis assay and then labeled with hashtag oligonucleotides (BioLegend) at 4°C for 30 minutes. Samples were then pooled and loaded into a Chromium Controller (10x Genomics) at a minimum concentration of 1,000 cells/µL, with a minimum target recovery goal of 20,000 cells. Library preparation was completed using the v3.1 Chromium Single Cell 3′ Kit (10x Genomics) and sequenced at Novogene on the NovaSeq X Plus with a target of 20,000 reads per cell and 5,000 reads per cell hashtag oligo-barcode library. The cell ranger software (version 7.2) was used to align samples to the rhesus macaque genome (Mmul_10), and subsequent analysis was carried out using Seurat (version 5.1). Briefly, the HTODemux function was used to assign identity. The cells with ambient RNA (<400 detected genes) or cells that were dying (>5% total mitochondrial gene expression) were removed. Cells from all samples were then integrated using Harmony, normalized using NormalizedData, and clustered using the first 10 principal components by the FindNeighbors and FindClusters Seurat functions (resolution=0.3). Canonical markers identified using the FindMarkers function were used to identify clusters and visualized by Uniform Manifold Approximation and projection (UMAP). DESeq (using default settings) determined differentially expressed genes (DEGs) in Seurat. Cell trajectories were reconstructed across pseudotime after dimensional reduction using Slingshot with a root state of C3. While the marker genes of C3 suggested that this cluster is in a differentiating state and is giving rise to other clusters, the expression of osteoclastogenic-related genes in this cluster was lower than in both the osteoclast (C5) and osteoclast precursors (C7), yet higher than in the remaining clusters. This indicates that C3 is positioned before C5 and C7. Gene ontology analysis of biological processes (GO-PB) was performed on the clusters’ marker genes using the PantherDB^27^. DEGs (FDR p-value ≤0.05, FC≥1.5) were enriched using the default settings of Reactome^28^.

### Conventional and spectral flow cytometry and dimensionality reduction analysis

Cells were incubated with antibodies targeting the cell surface in the presence of 50 µl of Brilliant Stain buffer (BD Biosciences), 5 µl of TruStain FcX Fc Receptor Blocking Solution (Biolegend), and 5 µl of True-Stain Monocyte Blocker (Biolegend) for 30 minutes at 4°C, protected from light. After incubation, the cells were washed with 1X PBS and subsequently permeabilized using Tonbo permeabilization solution (Tonbo Bioscience) for 2 hours at 4°C in the dark. The cells were washed using Tonbo Perm buffer and incubated with intracellular antibodies overnight at 4°C in the dark. Samples were acquired on a Cytek Aurora flow cytometer, which features 5 lasers (55 nm, 405 nm, 488 nm, 561 nm, and 640 nm) and 67 detectors and is operated with SpectroFlo Software v3.0. The cells underwent unmixing with stained beads or cells while enabling the autofluorescence extraction option. The resulting unmixed FCS files were analyzed using FlowJo v10.10.0. Undesired events were eliminated utilizing FlowAI 2.3.2^29^. The dimensionality reduction was performed using UMAP v4.1.1^30^. The clusters were identified by FlowSOM v4.1.0^31^ or Phenograph v2.5.0^32^. PHATE (Potential of Heat-diffusion for Affinity-based Trajectory Embedding; v1.0.0)^33^ was used to visualize cluster relations. To stain the cells for spectral flowcytometry, the following antibodies were used; CD3 (BUV496, OKT3, BD Biosciences), CD20 (BUV496, 2h7, BD Biosciences), CD14 (BUV805, MSE2, BD Biosciences), CD45RA (BUV395, 5H9, BD Biosciences), CX3CR1 (RB780, 2A9-1, BD Biosciences), NF-κB p65 (pS529) (PE, Kl 0-895.12.50, BD Biosciences), CD184 (CXCR4) (BV480, 12G5, BD Biosciences), HLA-DR (Spark Violet™ 538, L243, Biolegend), CD16 (APC/Fire™ 750, 3G8, Biolegend), CD34 (APC/Cyanine7, 561, Biolegend), CD282 (TLR2) (PE/Cyanine7, TL2.1, Biolegend), CD115 (CSF-1R) (PE/Dazzle™ 594, 9-4d21ef, Biolegend), SPI1 (PU.1) (Alexa Fluor® 647, 7C6B05, Biolegend), CD64 (VioBlue, 10.1.1, Miltenyi), TNF RI/TNFRSF1A (PE/Cy5.5, H398, Novusbio), CCR2 (APC, 48607, Novusbio), HIF-1 alpha (Alexa Fluor® 488, H1alpha67, Novusbio), CD38 (PerCP or PE/Cyanine7, AT1, Novusbio), CXCR1/IL- 8RA (Alexa Fluor® 700, 42709, Novusbio), CD123 (Brilliant Ultra Violet™ 737, 6H6, eBioscience), CD90 (Thy-1) (Brilliant Ultra Violet™ 563, eBio5E10 (5E10), eBioscience), CD127 (Super Bright™ 780, eBioRDR5, eBioscience), CD284 (TLR4) (Super Bright™ 600, HTA125, eBioscience), TNF alpha (Brilliant Violet™ 650, MAb11, eBioscience), IL1R2 (PE, 34141, Thermofisher), and IRF8 (PerCP-eFluor™ 710, V3GYWCH, eBioscience).

In order to assess the presence of osteoclast precursors, the bone marrow samples were stained with the following antibodies; HLA-DR (BV605, L243, Biolegend), CD123 (FITC, 6H6, Thermofisher Scientific), CD34 (APC-cy7, 561, Biolegend), CD115 (CSF1R) (PE, 9- 4D2-1E4, Biolegend), CD11b (PE-Cy7, CICRF44, Biolegend), TREM2 (Alexa Fluor™ 700, 237920, R&D Systems), RANK (Alexa Fluor™ 647, H-7, Santa Cruz), and CD14 (Pacific Blue, MSE2, BD Biosciences). Samples were acquired with the Attune NxT Flow Cytometer (Thermofisher Scientific, Waltham, MA). Supervised and unsupervised analysis was performed as described above.

### Colony-forming unit-granulocyte macrophage

The colony-forming unit-granulocyte macrophage (CFU-GM) assay was performed using MethoCult™ H4035 Optimum medium supplemented with recombinant human cytokines SCF, GM-CSF, IL-3, and G-CSF (StemCell) following manufacturer instructions. The CFU-GM assay mixture was prepared by mixing 0.3 ml of the 0.5×10^6^ cell in IMDM (Iscove’s Modified Dulbecco’s Medium; Gibco) with 2% FBS with 3 ml of MethoCult™ medium. To assess the impact of osteoclast-conditioned media on the generation of colonies, the control cells were mixed with 0.3 ml of osteoclast-conditioned media. An aliquot of the cell-medium mixture was dispensed into 35 mm culture dishes. The dishes were incubated at 37°C in a humidified 5% CO₂ atmosphere for 7 days. CFU-GM colonies were counted manually using a light microscope. To harvest the colonies, 1 ml of IMDM with 2% FBS was added to each culture dish. The methylcellulose medium was gently mixed by pipetting up and down, and the resulting cell suspension was transferred into a 15 mL centrifuge tube. The cell suspension was topped up to 10 mL with IMDM + 2% FBS. Samples were then centrifuged at 350×g for 10 minutes, and the cell pellet was carefully washed to remove residual methylcellulose. Cells were counted and stained with the following antibodies: CD45RA (V450, 5H9, BD Biosciences), CD3 (BV510, SP34, BD Biosciences), CD20 (BV510, 2h7, Biolegend), HLA-DR (BV605, L243, Biolegend), CD14 (BV711, MφP9, BD Biosciences), CD123 (FITC, 6H6, Thermofisher Scientific), CD34 (PE-Cy7, or APC-cy7, 561, Biolegend), CX3CR1 (APC, 2A9-1, Thermofisher Scientific), CD64 (APC or Alexa Fluor™ 700, 10.1.1, Thermofisher Scientific), and CD38 (PE-Cy7, AT1, Novusbio). The samples were acquired with the Attune NxT Flow Cytometer (Thermofisher Scientific). Supervised and unsupervised analysis was performed as described above.

### Luminex multiplex assay

The supernatants of differentiated osteoclasts were assessed for the presence of immune mediators using a non-human primate (NHP)-specific 5-plex (R&D Systems), including CCL2/MCP-1, IL-1β, IL-17a, IL-6, and PDGF-BB, following the manufacturer’s recommendations. Data was acquired using a MAGPIX instrument (Diasorin) and analyzed by a 5-parameter logistic regression with the xPONENT™ software (Diasorin).

### EdU cell proliferation assay

The proliferation assay was performed using Click-iT™ EdU (Thermofisher) following manufacturer instructions. Briefly, EdU (10 µM) was added to the culture of control cells that were incubated with or without osteoclast-conditioned media: RPMI: (1:1) for 6 hours. Cells were then collected and stained for cell surface markers, including HLA-DR (BV605, L243, Biolegend), CD14 (BV711, MφP9, BD Biosciences), CD123 (FITC, 6H6, Thermofisher Scientific), CD38 (PerCP, AT1, Novus), CD34 (PE-Cy7, 561, Biolegend), CX3CR1 (APC, 2A9-1, Thermofisher Scientific), CD64 (Alexa Fluor™ 700, 10.1.1, Thermofisher Scientific), and CD16 (APC/Fire™ 810, 3G8, Biolegend) for 30 minutes. The cells were then fixed with 4% paraformaldehyde for 15 minutes, permeabilized, resuspended in the reaction cocktail, including Pacific Blue™ fluorophore, and incubated for 30 minutes at room temperature, protected from light. Cells were then washed with PBS with 1% BSA, acquired with the Attune NxT Flow Cytometer, and analyzed using FlowJo.

### Statistical Analyses

We performed a normality assessment using the Shapiro-Wilk test with a significance level of alpha=0.05 and identified outliers using ROUT analysis at a Q value of 0.1%. If the data exhibited a normal distribution, we examined group differences through either paired or unpaired t-tests, applying Welch’s correction (parametric) or Mann-Whitney (non-parametric) as necessary. For multiple comparisons, we used the Holm-Sidak test, adjusting both family-wise significance and confidence levels to 0.05. For datasets that did not conform to Gaussian assumptions, we utilized the Mann-Whitney test for group comparisons. P-values of 0.05 or lower were considered statistically significant, while p-values between 0.05 and 0.1 were deemed as modest changes.

## RESULTS

### CAC enhances the capacity of HSPCs to differentiate into myeloid progenitor cells

To assess the impact of CAC on the differentiation potential of HSPCs to granulocyte-macrophage progenitors, we incubated bone marrow cells with SCF, G-CSF, GM-CSF, and IL-3. The colonies were imaged and quantified on day 7 post-incubation (**Fig. 1A**). A significant increase in the number of colonies was observed in samples derived from the CAC group (**Fig. 1B**). Colonies were then harvested and dissociated. The single cells were counted and phenotyped using flow cytometry (**Supp. Fig. 1A-B)**. It was observed that the absolute counts of CD14^-^CD34^+^ hematopoietic stem cells, common monocyte progenitors (cMoPs; Lin^-^CD38+CD34^+^CD123^+^CD45RA^+^CD64^+^), granulocyte-macrophage progenitors (GMPs; Lin^-^CD38+CD34^+^CD123^+^CD45RA^+^CD64^-^), and HLADR^+^CD14^+^ monocytes were significantly elevated in the CAC groups (**Supp. Fig. 1B and Supp. Fig. 2A**).

**Figure 1.**
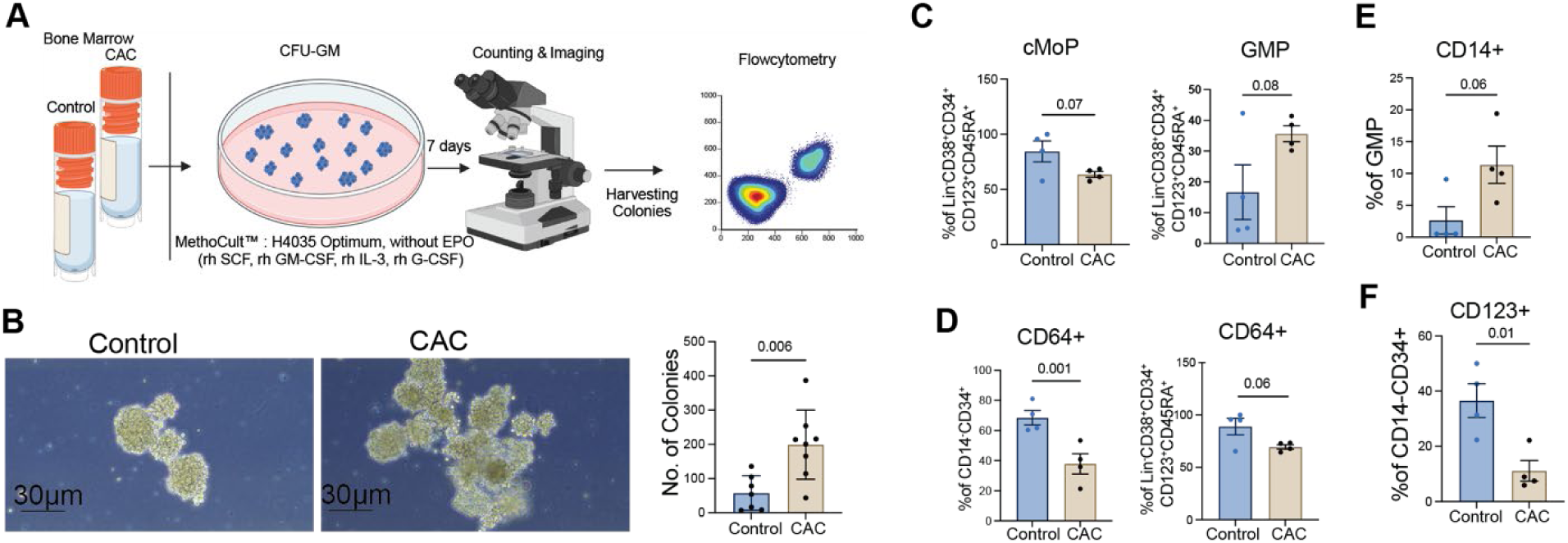
CAC increases the differentiation capacity of HSPCs into granulocyte-macrophage progenitors. **A**) Experimental design. **B**) Representative images of CFU-GM colonies and colony counts. **C**) The percentage of cMoP and GMP cells among their parent population. **D**) The percentage of CD64^+^ among CD14^-^CD34^+^ and Lin-CD38^+^CD34^+^CD123^+^CD45RA^+^. **E**) Percentage of CD14^+^ cells among GMPs and **F**) CD123^+^ cells among CD14^-^CD34^+^ cells. Error bars for all graphs are defined as ± standard error of the mean (SEM), and each dot corresponds to an individual animal.

We further evaluated the differentiation capacity of progenitor cells by calculating the percentage of descendant populations relative to their parent populations. A modest decrease in the frequency of cMoPs accompanied by a modest increase in GMPs within Lin-CD38^+^CD34^+^CD123^+^CD45RA^+^ was noted (**Fig. 1C**). This was in line with a significant decrease in CD64^+^ cells within the CD34^+^ progenitors as well as within Lin-CD38^+^CD34^+^CD123^+^CD45RA^+^, explaining the enhanced ratio of GMPs (CD64^-^) compared to cMoPs (CD64^+^) in CAC group (**Fig. 1D and Supp. Fig. 2B**). Further analysis revealed a modest elevation in the frequency of HLA-DR^+^CD14^+^ monocytes within the GMP population in CAC group (**Fig. 1E**). These data suggest that CAC skews differentiation of the Lin^-^CD38^+^CD34^+^CD123^+^CD45RA^+^ progenitor cells toward GMP-derived monocytes instead of cMoPs. These observations are in accordance with our prior observation that CAC is associated with the production of “neutrophil-like” inflammatory monocytes^34^. Indeed, a modest decrease in the frequency of CX3CR1^+^ cells among CD14^-^CD34^+^ and HLA-DR^-^CD14^+^ monocytes was observed (**Supp. Fig. 2C-D**). Reduced CX3CR1 expression on monocytes was shown to be indicative of a pro-inflammatory profile^35^.

Additional analysis showed a significant decrease in CD123^+^ cells among CD14^-^ CD34^+^ cell populations (**Fig. 1F**), which was maintained in downstream progenitors Lin^-^ CD38^+^CD34^+^CD123^+^CD45RA^+^ and cMoPs (**Supp. Fig. 2E**). CD123 is the receptor for IL- 3, playing a critical role in hematopoietic stem cell proliferation and differentiation^36^. Therefore, CD123 downregulation may drive the differentiation bias toward GMP rather than cMoP. Collectively, these findings support our previous observations^16^ and highlight that CAC results in preferential differentiation of hematopoietic stem cells towards granulocyte-monocyte lineage.

### CAC alters the abundance of HSCs and induces granulocyte-macrophage progenitors

To capture the impact of CAC on HSPCs, we analyzed bone marrow cells using spectral flow cytometry, incorporating 26 markers and employing dimensionality reduction techniques (gating strategy displayed in **Supp. Fig. 3A**). The supervised gating analysis revealed an increase in GMPs that was accompanied by a decrease in cMoPs within the Lin^-^CD38^+^CD34^+^CD123^+^CD45RA^+^ cells in CAC group (**Fig. 2A & Supp. Fig.3B)**. This observation corresponds with a reduction in the overall number of cMoPs (**Supp. Fig.3C**). Additionally, we detected a modest increase in common lymphoid progenitors (CLPs; Lin^-^ CD34^+^CD127^+^) in the CAC group^16^ (**Supp. Fig.3D**).

**Figure 2.**
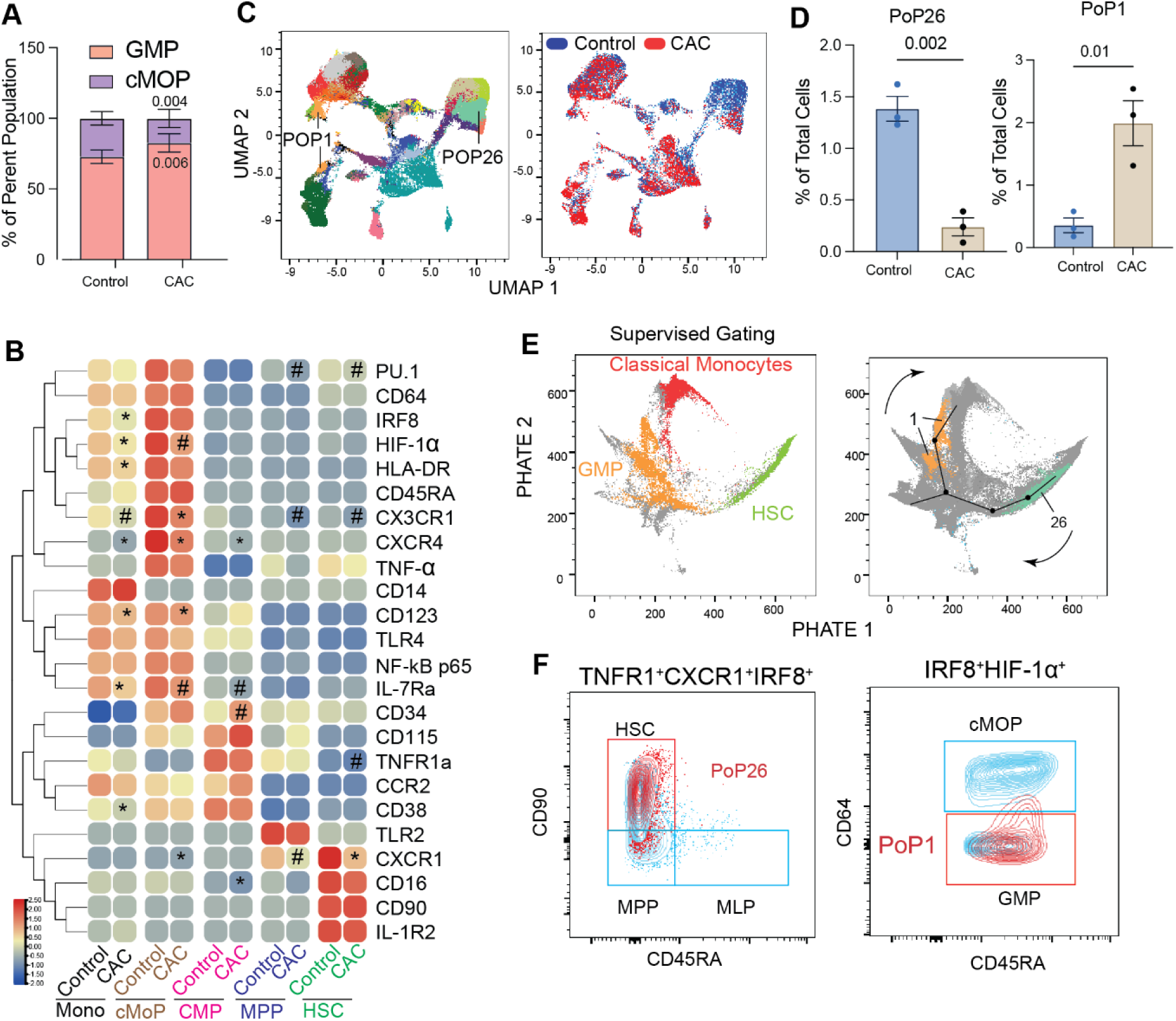
CAC depletes HSCs while enhancing GMPs. **A**) Frequencies of GMPs and cMoPs within Lin^-^CD38^+^CD34^+^CD123^+^CD45RA^+^ cells. **B)** Mean fluorescence intensity (MFI) of key hematopoietic markers; * indicates p≤0.05, and # indicates 0.05≤p≤0.1. **C**) UMAP depicts the clusters within the control and CAC groups. **D**) Contribution of control and CAC group in PoP1 and PoP26. **E**) The PoP1 and PoP26 of unsupervised analysis and monocytes, GMP, and HSC population of supervised analysis were mapped on the PHATE plot. Links between clusters were drawn manually. **F**) PoP1 and PoP26 overlapped on the gating strategy used for supervised analysis. Error bars for all graphs are defined as ± Standard deviation (SD).

We further evaluated the expression of regulatory cell surface proteins and key transcription factors within each population. Among the transcription factors tested, we observed a modest decrease in the expression of PU.1 in hematopoietic stem cells (HSCs; Lin^-^CD38^-^CD34^+^CD90^+^CD45RA^-^) and multipotent progenitors (MPPs; Lin^-^CD38^-^ CD34^+^CD90^-^CD45RA^-^) (**Fig. 2B).** Considering the function of PU.1, in myeloid lineage commitment^37^, this decrease could explain the reduction in cMoPs. The expression of HIF1-α was also downregulated in cMoPs and monocytes (**Fig. 2B)**. Furthermore, IRF8, as a transcription factor critical for myeloid lineage differentiation, was downregulated in mature monocytes in the CAC group (**Fig. 2B)**. Downregulation of IRF8 in monocytes increases the potential of their differentiation into osteoclasts^38^.

In line with our findings in the CFU-GM assay, the expression of CD123 was decreased in primary Lin-CD38^+^CD34^+^CD123^+^CD45RA^+^ cells (**Supp. Fig. 4A)**. In addition, the expression of CXCR1, CX3CR1, and CXCR4, which serve as receptors for IL-8, SDF-1 (CXCL12), and CX3CL1, respectively, was also decreased among various progenitors (**Fig. 2B**). These receptors are important for proliferation, cell migration, and inflammatory responses^39–41^. No differences were found in expression of CD64, CD45RA, CD14, TLR4, TLR2, NF-kb, and TNF-α, CD115, CCR2, CD90 and IL-1R2 (**Fig.2B)**.

We next conducted an unsupervised analysis to identify populations potentially overlooked in the supervised gating analysis. The FlowSOM algorithm identified 30 distinct clusters (**Supp. Fig.4B)**. Among them, PoP26 cluster was significantly decreased in the CAC group, while PoP1 was significantly enriched (**Fig.2C-D)**. To comprehensively characterize these clusters, we employed Marker Enrichment Modeling (MEM). PoP26 was composed of TNFR1^+^CXCR1^+^IRF8^+^ cells, indicating their higher potential response to TNF-α. PoP1 was identified as IRF8^+^HIF1-α^+^ cells. Higher levels of IRF8 in the PoP26 cluster could explain its depletion in the CAC group as a result of diminished self-renewal capacity^42^, and in PoP1, contributes to promoting the differentiation of lineage-committed progenitors to monocytes while suppressing neutrophil development^43^. HIF-1α boosts survival and proliferation in the relatively hypoxic bone marrow^44^. Therefore, the shift from Pop26 to Pop1 with CAC is aligned with the shift towards GMP-derived monocytes rather than cMOP.

Further, we used the PHATE analysis to yield additional insights into the developmental trajectories of the identified clusters. Mapping supervised gating on the PHATE graphs showed PoP26 overlaps with HSCs (Lin-CD38^-^CD34^+^CD90^+^CD45RA^-^) and PoP1 with granulocyte-monocyte progenitor (GMP; Lin^-^ CD38^+^CD34^+^CD123^+^CD45RA+CD64^-^) (**Fig.2E and Supp. Fig.4C).** Further overlapping PoP26 and PoP1 on supervised gating validated their contributions to HSCs and GMPs, respectively (**Fig.2F and Supp. Fig.4D)**. Collectively, all these data indicate that depletion of a TNFR1^+^CXCR1^+^IRF8^+^ HSC subset and enrichment of a IRF8^+^HIF1-α^+^ GMP subpopulation in the CAC group. These observations align with the findings of the CFU-GM assay and indicate that CAC skews hematopoietic stem cells toward the enrichment of granulocyte-monocyte progenitors, serving as potential precursors for osteoclasts.

### CAC increased osteoclast size and number of nuclei per osteoclast

To investigate whether CAC’s mediated increase in the frequency of myeloid progenitors leads to an increase in osteoclast precursors, we incubated total bone marrow cells with RANKL and M-CSF. The differentiated cells were stained for tartrate-resistant acid phosphatase (TRAP). Additionally, to assess the bone resorption activity, the cells were cultured on plates coated with calcium phosphate bound to fluorescein amine-labeled chondroitin sulfate, which is released following osteoclast resorption activity (**Fig. 3A**). Our results indicated that the total number of osteoclasts - defined as TRAP-positive cells with three or more nuclei - was significantly higher in the CAC group compared to the control (**Fig. 3B and Supp. Fig. 5A**). We further stratified the osteoclasts by size and noted a significant increase in the number of giant cells (>10,000 μm²) in the CAC group (**Fig. 3C**). Additionally, the osteoclast area per well (**Supp. Fig. 5B**) and the osteoclast area per cell-cell surface area- (**Supp. Fig. 5C**) were also significantly enhanced in the CAC group. Moreover, on average, 50% of those derived from CAC had over ten nuclei (**Fig. 3D**). Finally, growing the cells on a calcium-coated plate demonstrated an increase in the pit area in the CAC group (**Fig. 3E**) and an increase in fluorescence intensity of fluorescein amine-labeled chondroitin sulfate released from calcium phosphate layer in the CAC group on day 5 (**Supp. Fig. 5D**).

**Figure 3.**
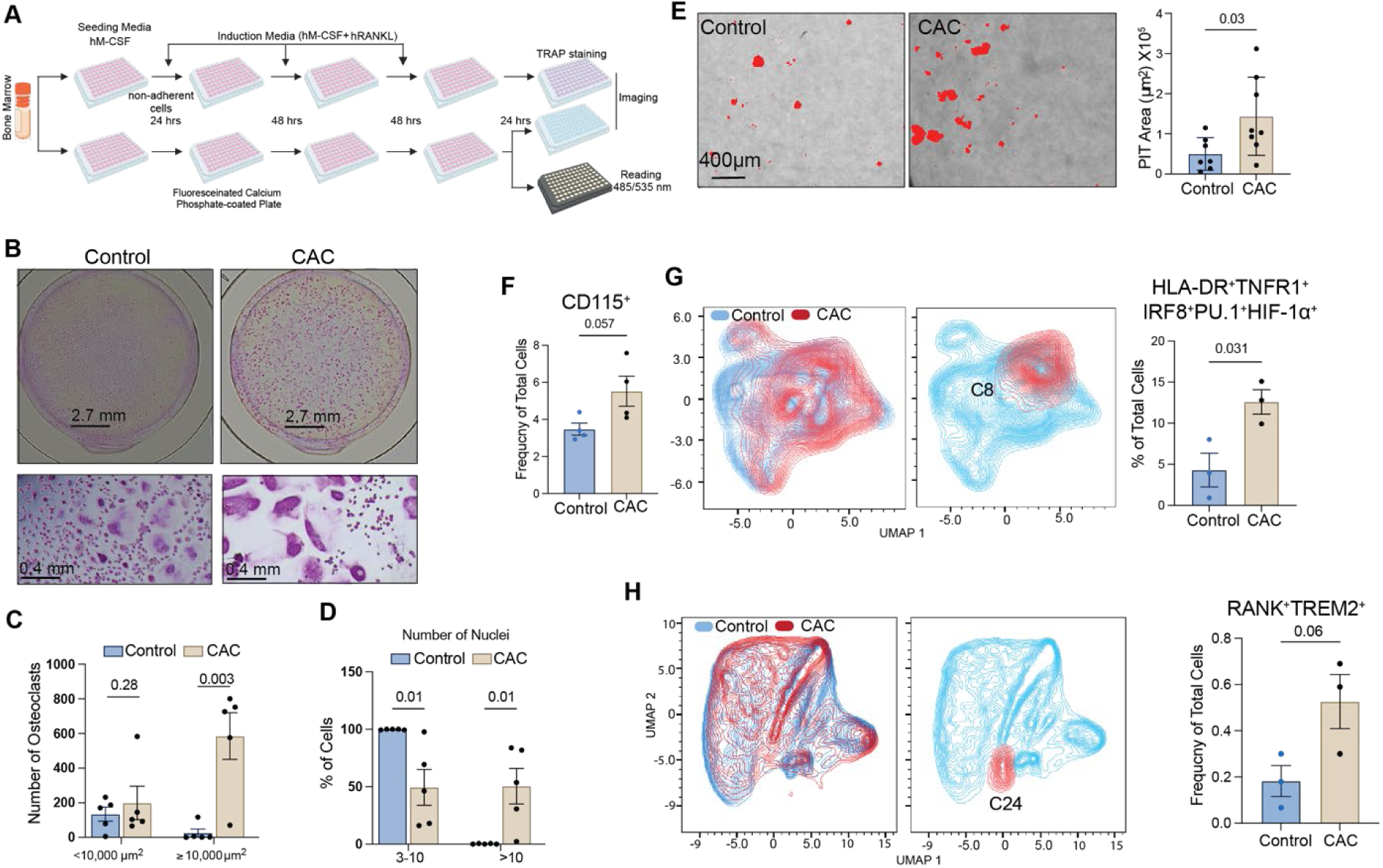
CAC enhances the number and size of osteoclasts. **A)** Experimental design to assess the impact of CAC on osteoclastogenesis. **B**) Representative images of differentiated osteoclasts are provided at 4X and 10X. Osteoclasts were classified according to their **C**) size and **D**) number of nuclei per cell. **E**) Representative images (4X) of the pit area following differentiation of bone marrow cells on calcium-coated plates. **F**) The frequency of CD115^+^ bone marrow cells in the experiment presented in Fig.2. **G**) Dimensional reduction analysis of the CD115^+^CD14^+^ population showing the frequency of C8. **H**) Dimensional reduction analysis showing the frequency of Cluster 24 (RANK^+^TREM2^+^). Data represent mean values ± SEM.

The increased frequency and size of osteoclasts combined with increased bone resorption activity observed in the CAC group suggest that CAC may lead to an increased frequency of osteoclast precursors. Since CD115 (CSF1R) is continuously expressed on common-myeloid progenitors (CMPs) to committed pre-osteoclasts^45–48^, we assessed the impact of CAC on its expression. Frequency of CD115^+^ cells was modestly higher in CAC samples (**Fig. 3F).** Since osteoclast precursors express CD115 and CD14, we extracted this population to conduct dimensional reduction analysis. Among the 13 identified clusters, we observed a significant increase in cluster 8 (C8; HLA-DR^+^TNFR1^+^IRF8^+^PU.1^+^HIF-1a^+^) within the CAC samples (**Fig. 3G & Supp. Fig. 5E**). Elevated levels of both IRF8 and PU.1 suggest that this cluster could represent a transitional stage between osteoclast precursors and fully committed pre-osteoclasts.

To assess the committed osteoclast precursors, we designed an additional staining panel that included RANK. Dimensional reduction and clustering analysis on CD115^+^ cells identified 25 clusters, among which 7 were RANK^+^ (**Supp. Fig. 5F)**. A modest increase in cluster 24 (RANK^+^TREM2^+^) (p=0.06) **(Fig. 3H-I)** was observed in the CAC group. TREM2 plays a critical role in proliferation and Ca2^+^ mobilization^49–51^. Together, these observations suggest that the enhanced ability of CAC-derived bone marrow cells to differentiate into osteoclasts is correlated to an increase in osteoclast precursors in the bone marrow.

### CAC alters the transcriptome profile of osteoclast precursors

To further assess the impact of CAC on osteoclast differentiation, we collected the *in vitro* differentiated cells and conducted scRNA-seq analysis. The subsequent UMAP clustering of a total of 15,445 cells revealed seven distinct cellular states (**Fig. 4A**). Cluster 1 was classified as “terminally differentiated macrophages,” characterized by elevated levels of macrophage-related genes such as *CD14*, *MAMU-DRA* (HLA-DR), *CD36, MMP2*, *CD74*, *C5AR1*, *C1QC*, *VSIG4*, and *SIGLEC1* (**Fig. 4B & Supp. Fig. 6A**). Cluster 2 was classified as “intermediate macrophages” displaying high levels of *CD14* and *MAMU-DRA*, alongside significant expression of *MX1*, *SAMHD1*, *IFT1* (**Fig. 4B**), *STAT2*, *PDE4B*, and *CCND1* (**Supp. Fig. 6A**). The marker genes of clusters 1 and 2 play roles in macrophage functions, including phagocytosis, antigen processing and presentation, and macrophage activation (**Supp. Fig. 6B**). Cluster 4 was identified as “dendritic-cell like macrophages” based on elevated expression of *CD52* and *ITGAX* (*CD11c*) in addition to macrophage markers *CD14* and *MAMU-DRA* (**Fig. 4B & Supp. Fig. 6A**). Cluster 6 displays a high expression of canonical macrophage markers *CD14* and *MAMU-DRA*, along with increased levels of cell cycle-related genes *TOP2A*, *MIK67*, *CCNB1, CENPE,* and *NUSAP1* (**Fig. 4B & Supp. Fig. 6A**). The marker genes for this cluster enriched for “G2/M phase transition” and “regulation of cell cycle checkpoint”, leading to the designation of cluster 6 as “proliferating macrophages” (**Supp. Fig. 6B**).

**Figure 4.**
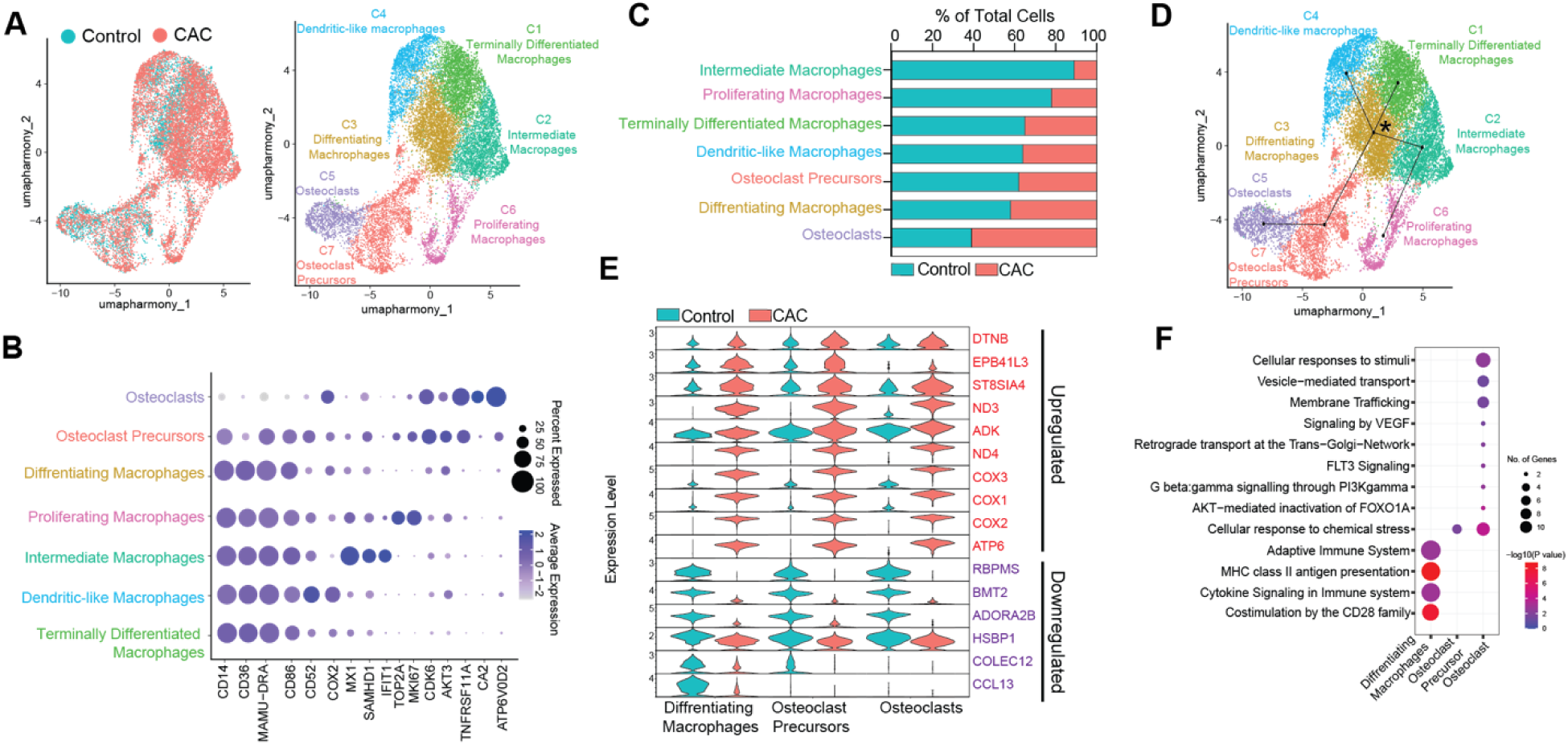
Single-cell RNA-sequencing reveals CAC-mediated transcriptome changes in osteoclasts. **A)** UMAP after dimension reduction of 15,445 cells representing control and CAC cells and identified clusters. **B**) A shortlist of gene markers that was utilized to identify clusters. **C**) Percentage of each group contributing to each cluster. **D**) UMAP clustering with Slingshot lineage projection lines. **E**) Expression levels of selected DEGs in specified clusters. **F**) Functional enrichment of genes upregulated with CAC in “differentiating macrophage”, “osteoclast precursor”, and osteoclast” clusters using Reactome. The size of the bubbles represents the number of genes associated with each GO term, and the bubble color indicates the statistical significance of the enrichment of a GO term based on the -log10 of the p-value.

Cluster 3 was classified as “differentiating macrophages” given expression of macrophage markers (*CD14*, *MAMU-DRA*, and *CD36*) as well as high levels of *CDK6* and *AKT3* (**Fig. 4B & Supp. Fig. 6A)**. The genes in this cluster enriched to “positive regulation of cell differentiation” and “actin cytoskeleton organization” (**Supp. Fig. 6B**). Cluster 7 was defined as “committed osteoclast precursors” based on the high expression levels of *CDK6* and *AKT3*, coupled with the expression of osteoclast-related genes such as *TNFRSF11A*, *OSCAR*, and *SLC7A11* (**Fig. 4B, Supp. Fig. 6A**). The genes in this cluster mapped to “actin cytoskeletal organization”, “cell differentiation,” and “osteoclast differentiation,” suggesting that these cells are undergoing major structural changes to differentiate into osteoclasts (**Supp. Fig. 6B**). Finally, Cluster 5 was identified as “terminally differentiated osteoclasts,” marked by a high expression of osteoclast-specific markers, including *TNFRSF11A*, *CA2*, and *ATP6V0D2* (**Fig. 4B**) as well as *NFATC1*, *MMP9*, *RAC2*, *DCSTAMP*, *JPD2*, and *ACP5* (**Fig. 4B, Supp. Fig. 6A**). The genes in this cluster mapped to terms including “bone resorption” and “proton transmembrane transport” (**Supp. Fig. 6B**).

The analysis of the proportional representation of these clusters within the total cell populations revealed that approximately 60% of the osteoclast population is comprised of cells from the CAC samples (**Fig. 4C, Supp. Fig. 6C**). We conducted a trajectory analysis to better comprehend the relationships among these clusters. The clusters were organized by pseudo-time, beginning with “differentiating macrophages” (C3) and progressing to “terminally differentiated macrophages (C1), intermediate macrophages (C2), and dendritic-like macrophages (C4). “Intermediate macrophages” (C2) further gave rise to “proliferating macrophages” (C6), and the “differentiating macrophage” cluster transitioned into an “osteoclast precursor” cluster (C7), which ultimately developed into “osteoclasts” (C5) (**Fig. 4D)**

Next, we assessed the expression of genes involved in osteoclastogenesis within all clusters. An enhanced expression of these genes was observed in “differentiating macrophages” (C3), “osteoclast precursors” (C7), and “osteoclasts” (C5) (**Supp. Fig. 7A)**. Interestingly, in “differentiating macrophages” expression of kinases including *MAKP1*, *SYK*, and transcription factors such as *SPl1*, *NFKBIA*, *RELB*, *MITF*, and *REST* was increased indicating enhanced transcription machinery. Additionally, in the “osteoclast precursor” cluster, expression of genes involved in structural (*FYN*, *TNFRSF11A*, *DOCK5*, *FLNA)* and metabolic processes (*ITPR1*, *S100A2, SLC7A11)* was also increased, indicating commitment towards terminally differentiated osteoclast. In the “osteoclast” cluster, we noted a marked expression of key osteoclast genes, such as *SEMA4D*, *NFATC1*, *RAC2*, *BMPR1A*, *CA2*, *CALCR*, *SLC4A2*, and *ACP5*. (**Supp. Fig. 7A)**.

We next extracted the DEGs between the CAC and control groups within these three clusters. Significantly higher expression of *ND3*, *COX3*, *COX1*, *COX2*, *ATP6*, and *ND4* was detected in the CAC cells compared to the control group. (**Fig. E**). In the “differentiating macrophage” cluster, we detected a significant increase in the expression of genes important for osteoclastogenesis, such as *NAMPT* and *SLC7A11*. The upregulated genes in the “differentiating macrophage” cluster were primarily associated with important macrophage signaling pathways, including “cytokine signaling,” the “adaptive immune system,” “MHC class II antigen presentation,” and “costimulation via CD28” (**Fig. 4F & Supp. Fig. 7B).**

Expression of ATP-related genes, including *ATP8B4* and *AGTPBP1,* as well as the calcium regulator *SLC8A1,* was significantly increased in the CAC-derived “osteoclast precursor” cluster compared to the control (**Supp. Fig. 7B)**. The genes upregulated with CAC in the “osteoclast precursor” cluster enriched for the “cellular response to chemical stress” (**Fig. 4F)**. Furthermore, expression of genes important for the “AKT-mediated inactivation of FOXO1a”, “stimulated vesicle-mediated transport”, and “membrane trafficking processes” (**Fig. 4F**) such as *SLC18B (a* protein transporter), *RALBP1* (involved in receptor-mediated endocytosis), and *AKT3* were significantly increased within the CAC-derived “osteoclast” cluster compared to controls (**Supp. Fig. 7B)**. Finally, negative regulators of mTOR and NF-kb signaling such as *BMT2* and *ADORA2B* were downregulated in all three clusters with CAC. These data, combined with previously presented results, indicate that CAC impacts key signaling pathways involved in osteoclast differentiation. This could elevate the risk of osteoclast generation after the incidence of a pathological disease following CAC.

### CAC-induced osteoclastogenesis alters myeloid production

Although osteoclasts are dispensable for the maintenance and mobilization of hematopoietic stem cells^52^, enhanced osteoclastogenesis may impact hematopoiesis via altered release of immune mediators^51^. To evaluate this hypothesis, we conducted a CFU-GM assay using conditioned media obtained from osteoclasts of both control and CAC groups on day 5 (**Fig. 5A**). We detected an enhanced number of colonies formed in cultures supplemented with CAC-derived conditioned media (**Fig. 5B**). We collected these colonies and evaluated them using flow cytometry analysis. A modest increase was observed in the frequency of granulocyte-macrophage progenitor Lin^-^ CD38^+^CD34^+^CD123+CD45RA^+^among the CD34^+^CD38^+^ population (**Fig. 5C**).

**Figure 5.**
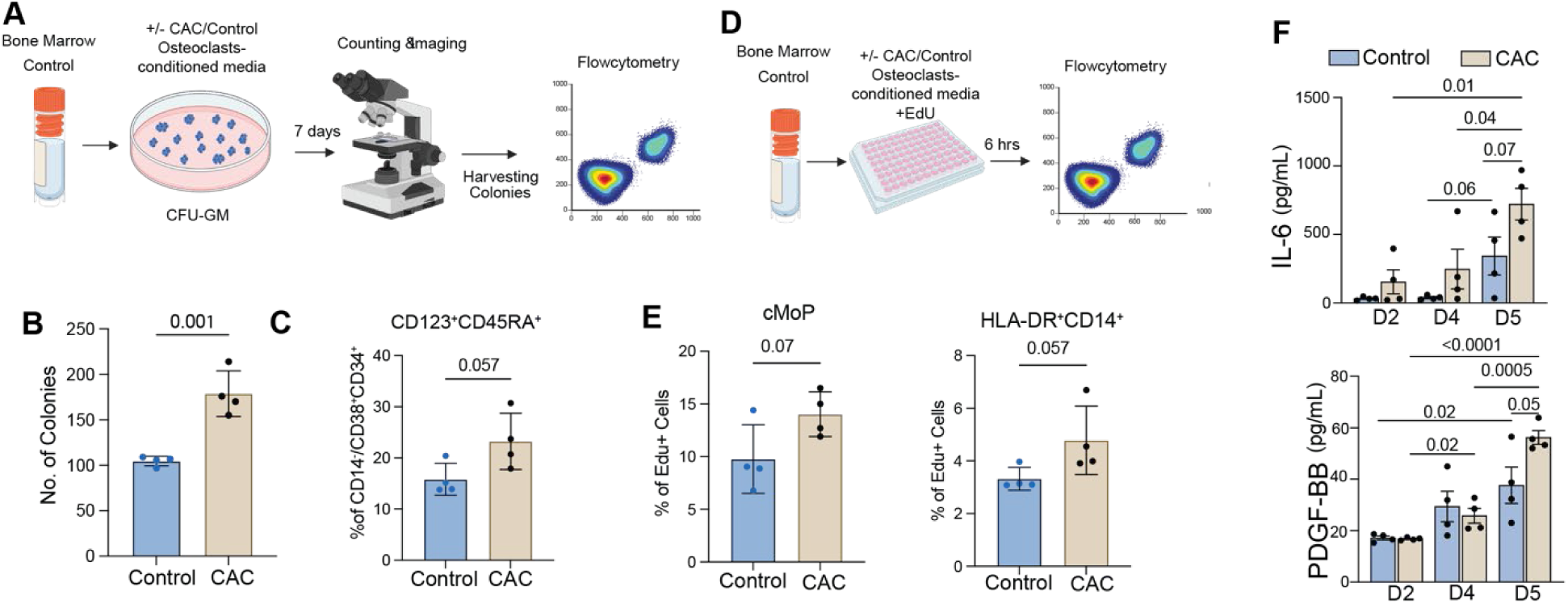
Osteoclastogenesis induced by CAC impacts myeloid production. **A)** Experimental design used to perform the CFU-GM assay using conditioned media obtained from osteoclast cultures of control and CAC groups. **B)** The number of CFU-GM colonies. **C)** The ratio of Lin^-^CD38^+^CD34^+^CD123+CD45RA^+^ among the CD34^+^CD38^+^ in CFU-GM assay. **D**) Experimental design to perform EdU assay. **E**) The frequency of cMoP EdU^+^ and HLADR^+^CD14^+^EdU^+^ in cultures incubated with osteoclast-conditioned media. **F**) Levels of IL-6 and PDGF-BB measured in supernatants of the osteoclasts culture. Data represent mean values ± SEM

Further, to assess the impact of osteoclast-conditioned media on the proliferation rate of myeloid progenitors, we incubated control bone marrow cells with control or CAC-derived osteoclast-conditioned media in the presence of EdU (**Fig. 5D**). A modest increase in the population of cMoP EdU^+^ and HLADR^+^CD14^+^EdU^+^ cells was observed in CAC group (**Fig. 5E**). To identify factors that promote osteoclast differentiation, we collected the supernatant of cells differentiating into osteoclasts on days 2, 4, and 5. This supernatant was analyzed for the presence of factors known to be released by osteoclasts, including CCL2/MCP-1, PDGF-BB, IL-1β, IL-17A, and IL-6. We observed an increase in the release of IL-6, PDGF-BB (**Fig. 5F**), and MCP-1 (**Supp. Fig. 8**) in the CAC group. These findings indicate that CAC promotes both the differentiation of osteoclast precursors and their immune mediator production.

## DISCUSSION

Bone turnover is a continuous physiological process involving both resorption and synthesis. Osteoporosis occurs when there is a disruption between bone breakdown and formation^53^. While primary osteoporosis is bone loss caused by normal aging or postmenopausal status, secondary osteoporosis is characterized by low bone mass and microarchitectural changes in the bone associated with underlying diseases or medications^54–56^. CAC may directly cause secondary osteoporosis^57, 58^ as well as exacerbate bone loss in the context of underlying diseases or medication^59, 60^ through dysregulation in the activity of osteoblasts^61^ and osteoclasts^62^.

Prior studies using a macaque model of voluntary ethanol self-administration noted that as little as 3 months of ethanol consumption decreased bone turnover^20^. Six months of voluntary alcohol consumption in male cynomolgus macaques was reported to reduce intracortical bone porosity^19, 21^. Following twelve months of CAC in male rhesus macaques, cancellous bone formation in the lumbar vertebra was reduced^17^, and suppression in intracortical bone remodeling was reported, as evidenced by lower cortical porosity and labeled osteon density^18^. Nevertheless, further cellular and molecular research is needed to investigate the mechanism underlying CAC-mediated bone remodeling in this model.

We evaluated the ability of bone marrow cells to differentiate into granulocyte-monocyte progenitors as potential precursors for osteoclasts. We discovered that the enhanced number of CFU-GM colonies derived from CAC bone marrow cells aligned with a greater ability of the progenitors to develop into GMPs instead of cMoP. This finding is consistent with studies from our group and others that showed that alcohol intake is linked to the disruption of hematopoiesis, particularly impacting myeloid progenitor cells^16, 63–65^. This myeloid differentiation bias may be a result of DNA lesions in HSCs caused by acetaldehyde^66^. The imbalance of cMoP/GMP may also be mediated by the downregulation of CD123 (IL-3 receptor) on Lin-CD38+CD34+CD123+CD45RA+ cells, given IL-3’s role in the proliferation and differentiation of HSCs^36^. Additionally, IRF8 and HIF-1α expression was increased with the CAC group, which could explain the increase in GMP cell numbers, as bone marrow is generally more hypoxic than other tissues, making progenitors with high *HIF1A* expression more prone to proliferation and differentiation^44^.

Further characterization of hematopoietic stem and progenitor cells showed a modest decrease in PU.1 level in HSCs and significant depletion of an HSC subpopulation (TNFR1^+^CXCR1^+^IRF8^+^) in the CAC group. Reduced HSCs can occur due to several factors, such as diminished self-renewal or increased differentiation and migration. Previous studies have shown that elevated RANKL promotes HSC mobilization^67^. Additionally, considering high TNF-α receptor expression in these cells, TNF-α may drive myeloid regeneration^68^. Moreover, PU.1 plays a critical role in lymph-myeloid lineage commitment^37^ and its constitutive expression in HSCs is necessary to maintain the HSC pool in the bone marrow^37^. Therefore, decreased expression of PU.1 could indicate accelerated differentiation of HSC.

The equilibrium among lineage-restricted progenitors, such as cMoPs and GMPs, is crucial for maintaining the balance of monocyte/macrophage-derived cells, including osteoclast precursors. Therefore, to assess if an enhanced GMP population may lead to increased osteoclast, we performed *in vitro* osteoclastogenesis. We observed a rise in TRAP-positive cells within the CAC group, which was associated with both larger cell size and greater nuclear count. Osteoclasts are characterized by their multinucleated structure. The number of nuclei in each osteoclast determines the extent to which osteoclasts can resorb a bone^69^. As a result, we noted a significant increase in the resorption area in the CAC group. Furthermore, a greater number of nuclei suggests an enhanced potential for pre-osteoclasts to fuse and form multinucleated cells^69^.

We then assessed pre-osteoclasts in primary bone marrow cells through flow cytometric phenotyping. Dimensional reduction analysis revealed that HLA-DR^+^TNFR1^+^IRF8+PU.1^+^HIF-1α^+^ within CD115^+^ cells could serve as potential osteoclast precursors. However, by including RANK in the panel, we found that a cluster of positive for both RANK and TREM2 enriched in the CAC group. TREM-2 binds with DAP12, resulting in the tyrosine phosphorylation of DAP12- ITAM facilitates Ca2^+^ mobilization, activation of PKC and MAP kinases, and the restructuring of the actin cytoskeleton^49–51^. In addition, the nuclear translocation of various transcription factors, triggered by signals from TREM2 and RANK, leads to the proliferation and differentiation of pre-osteoclasts, resulting in the transcription of essential osteoclast genes like *DC-STAMP*, *MMP9*, *CTSK*, and *ACP5*, culminating in forming a mature osteoclast^70^.

While *in vitro* and *in vivo* experiments have uncovered multiple signaling pathways involved in osteoclastogenesis, recent studies utilizing scRNA-seq of primary bone marrow cells^71, 72^ and *in vitro* differentiated cells^73^ have provided deeper insights into these processes by improving our understanding of the heterogeneity within osteoclast precursors, revealing distinct cellular states and regulatory mechanisms that drive their differentiation. We analyzed scRNA-seq data from *in vitro* differentiated cells and captured multiple cell states ranging from macrophages to differentiated osteoclasts. Our data are aligned with previous studies showing the monocyte precursors go through a series of processes, including membrane raft assembly, proliferation and RNA metabolism, energy metabolism, and differentiation into mature osteoclasts^73, 74^.

When assessing the DEGs between CAC and control, we found a dysregulation of genes that regulate NF-κB and ROS production, as well as suppressing pro-inflammatory cytokines across all three osteoclast clusters. We also detected enhanced expression of genes, leading to increased mitochondrial ATP production and proliferation. The upregulation of COX proteins enhances the conversion of arachidonic acid into prostaglandins, which increases RANKL production, reduces OPG-a decoy receptor that binds to RANKL- and influences osteoclast precursor survival and differentiation via cAMP and protein kinase A (PKA)-mediated signaling pathways^75–77^.

In the “differentiating macrophage” cluster, we noted a significant increase in the expression of genes previously associated with osteoclastogenesis, such as *NAMPT* and *SLC7A11* with CAC. NAMPT (Nicotinamide phosphoribosyltransferase) plays a role in NAD synthesis. Further, NAMPT expression, by inhibiting histone acetylation while enhancing histone methylation of the osteoclast key protein *NFATc1* promoter, enhances its expression and promotes osteoclastogenesis^78^. NFATc1 will promote the transcription of *SLC7A11,* which plays a role in importing oxidized cysteine. Interestingly, inhibition of cysteine reduction was shown to reduce osteoclastogenesis by selectively targeting osteoclast precursors^79^.

In the “osteoclast precursor” cluster, we observed a significant increase in the expression of membrane transporters with CAC-derived *ATP8B4* and *SLC8A1*. *ATP8B4 (*ATPase Phospholipid Transporting 8B4) is involved in phospholipid transport in the cell membrane. However, there has not been much study on the functional role of this protein. However, its role may be involved in vesicular trafficking between the Golgi and plasma membrane^80^. The Golgi apparatus is a prominent organelle in osteoclasts. Additionally*, SLC8A1* (Solute carrier family 8 member A1) mediates the exchange of Ca^2+^ and Na^+^ across the cell membrane and plays an accessory role in osteoclast Ca^2+^ homeostasis^81^. In the CAC-“osteoclast” cluster, we noticed upregulation of genes, including *SLC18B and RALBP1.* SLC18B belongs to the solute carrier (SLC) group of membrane transport proteins and actively transports polyamines, including spermine and spermidine, through the exchange of H^+82^. However, the role of *SCL178B* is not known in osteoclastogenesis, as polyamines play a role in numerous cellular functions, including cell growth, differentiation, and survival; SCL178B may enhance osteoclast activity. Additionally, the enhancement of spermidine/spermine has been associated with osteoclastogenesis^83^. RALBP1 plays a role in receptor-mediated endocytosis and may contribute to the bone resorption activity of osteoclasts. Collectively, these genes suggest significant cell membrane activity in osteoclasts derived from CAC.

Bone niche maintains the self-renewal ability, multipotency, and quiescence of HSCs^84^. Therefore, secretions by osteoclasts may disrupt the HSCPs’ hemostatic differentiation. We found that CAC-derived osteoclast-conditioned media led to increased GMPs. Given that, among all clusters, the frequency of the “osteoclast” cluster frequency in CAC groups was higher than in control, osteoclasts likely produce IL-6 and PDGF-BB. Nevertheless, experiments, such as intracellular cytokine staining, are needed to identify the sources of these cytokines. Regardless of the source, the impact of osteoclast-conditioned media was significant on myeloid cell production. IL-6 was shown to play a role in osteoclastogenesis^62^ and has also been implicated in regulating hematopoiesis^85–87^. Therefore, in an inflammatory condition, not only osteoclast precursors may give rise to osteoclasts^88^, but they may also impact hematopoiesis. However, the interaction between osteoclasts and HSC may involve more than merely cytokine effects. The elevated calcium levels resulting from bone resorption in this microenvironment play a role in HSC maintenance as they express seven membrane-spanning calcium-sensing receptors (CaR)^89^.

In summary, data presented here show that alcohol intake enhances hematopoietic stem cells’ capacity to differentiate into osteoclast precursors. These precursors can mature into osteoclasts when osteoclastogenic signaling is activated. Further research is necessary to explore the molecular mechanisms driven by alcohol that promote the generation of osteoclast precursors. Uncovering these mechanisms will help us identify treatment strategies to combat alcohol-related reduced bone density and fracture.

## AUTHOR CONTRIBUTIONS

HH performed experiments and analyzed the data with the help of MB. MB prepared the RNA-seq library. MB and H.H. analyzed the RNA-seq data. HH wrote the manuscript with MB and IM’s assistance. JK assisted in osteoclast counting. KG designed and oversaw the live alcohol self-administration studies, RG helped implement the self-administration studies, oversaw the necropsy procedures, and provided the bone marrow samples through the MATRR.com research resource. IM reviewed and edited the manuscripts. All authors have read and approved the final draft of the manuscript.

## ACKNOWLEDGMENTS

We thank members of the Messaoudi laboratory for their help and feedback. We thank Dr. Delphine Malherbe for reading the manuscript and providing feedback. This work was supported by NIH 1R01AA028735-01 (I.M.), 5U01AA013510-20 (K.A.G.), 2R24AA019431-11 (K.A.G.), P-51OD011092, and Pilot fund from the University of Kentucky’s Substance Use Priority Research Area (SUPRA) supported by the Vice President for Research (HH).

## Data and code availability

Additional information and requests for resources should be directed to Dr. Ilhem Messaoudi. The datasets supporting the conclusions of this article are available on NCBI’s Sequence Read Archive (SRA# pending).

---

**Supp. Fig.1.**
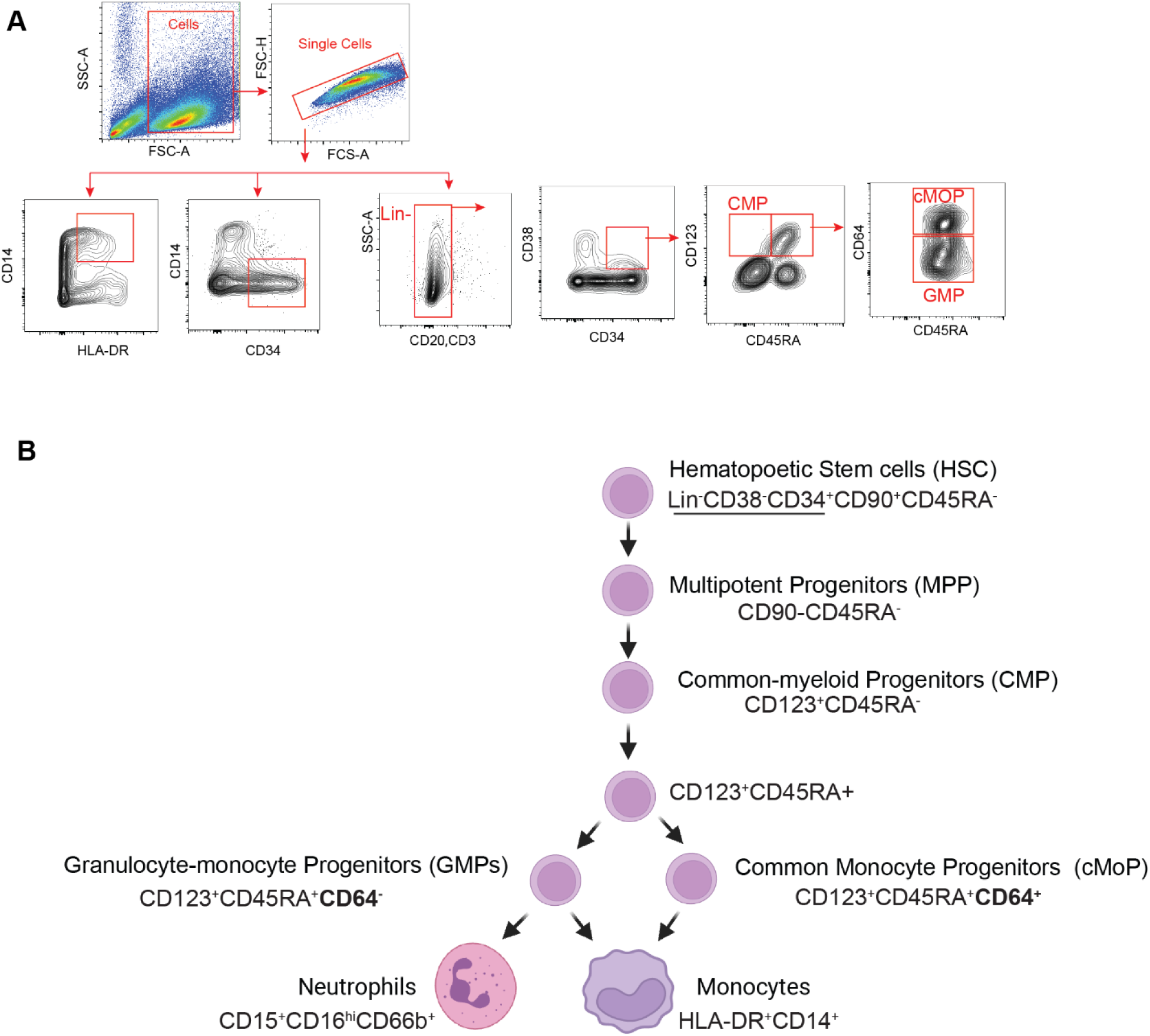
CAC skews HSPCs differentiation to myeloid cells. **A)** Gating strategy. **B)** An overview of monopoiesis in the bone marrow of rhesus macaques excluding monocyte-dendritic progenitors.

**Supp. Fig.2.**
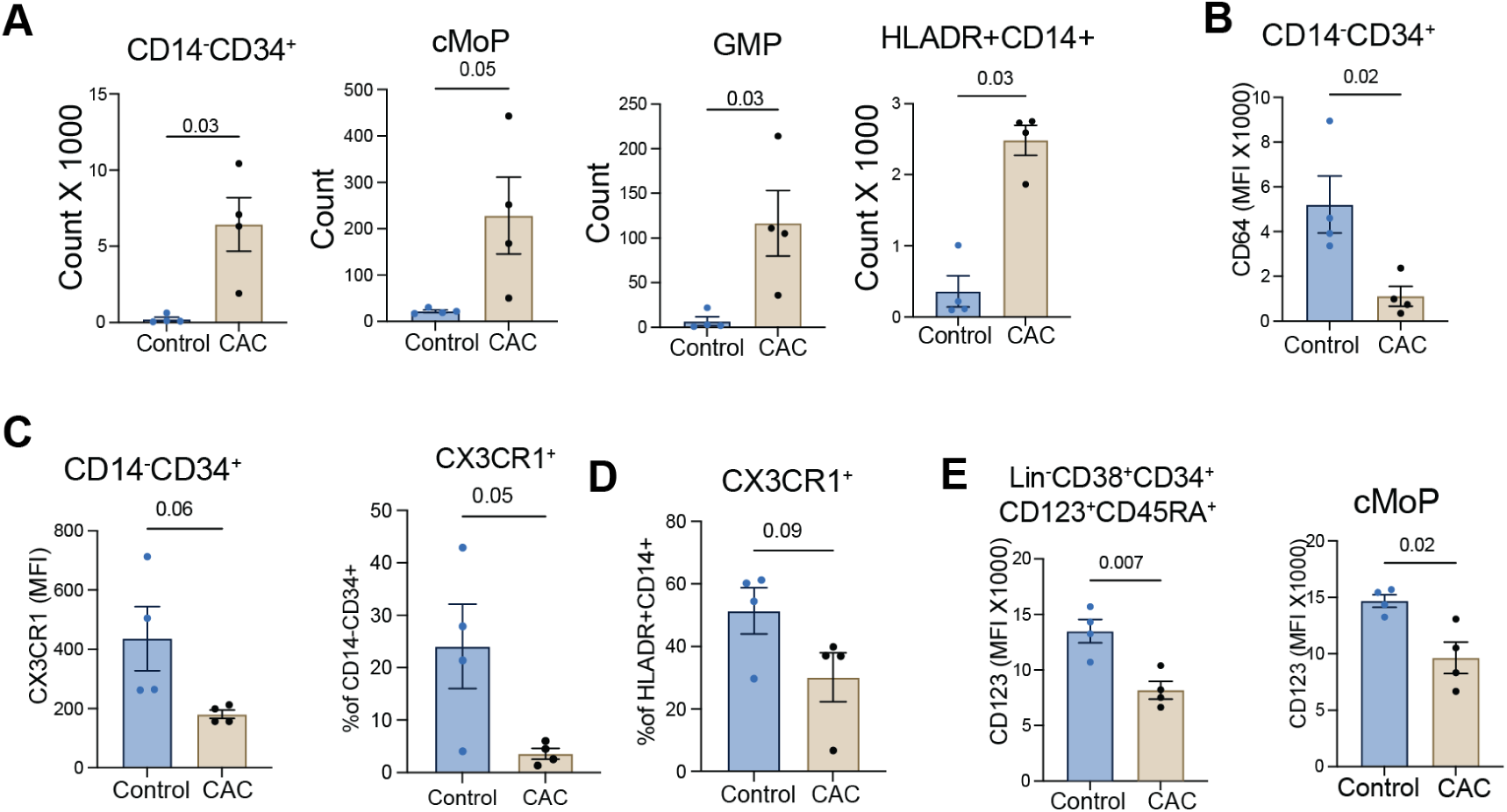
CAC skews HSPCs differentiation to myeloid cells. **A)** Absolute counts of each population following harvesting colonies and identifying populations by flow cytometry. **B**) MFI of CD64 on CD14^-^CD34^+^ cells. **C)** MFI of CX3CR1 on CD14^-^CD34^+^ and the percentage of CD14^-^CD34^+^CX3CR1^+^. **D**) The percentage of HLA-DR^+^CD14^+^CX3CR1^+^ cells. **E)** MFI of CD123 on Lin^-^CD38^+^CD34^+^CD123^+^CD45RA^+^ and cMoP cells.

**Supp. Fig.3.**
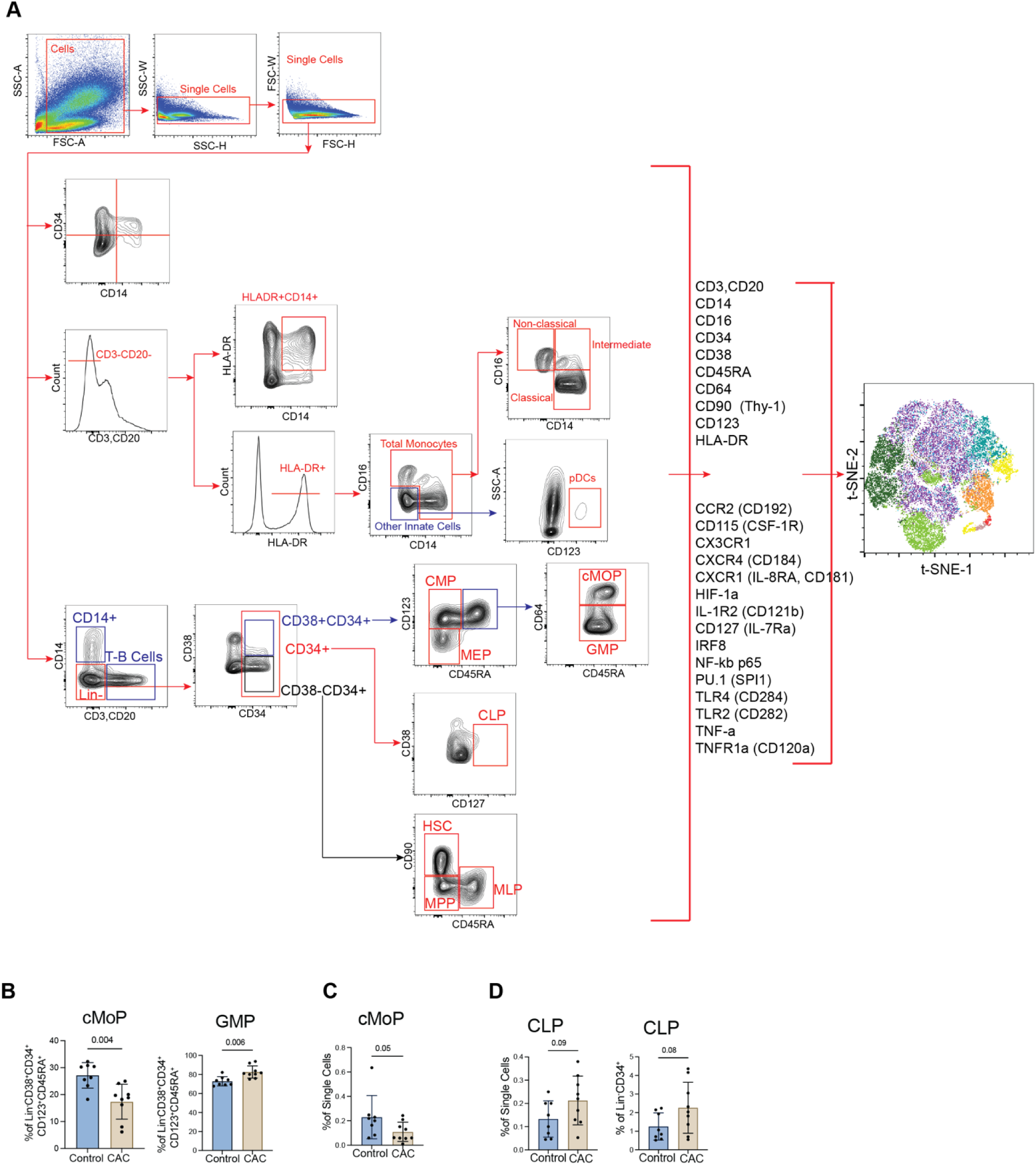
CAC dysregulates HSPCs differentiation. **A**) The gating strategy used for spectral flow cytometry analysis. **B**) Differentiation of Lin^-^CD38^+^CD34^+^CD123^+^CD45RA^+^ cells into cMoPs and GMP. **C**) The percentage of cMoP among all bone marrow cells. **D**) The percentage of CLPs among all bone marrow cells and their parent population.

**Supp. Fig.4.**
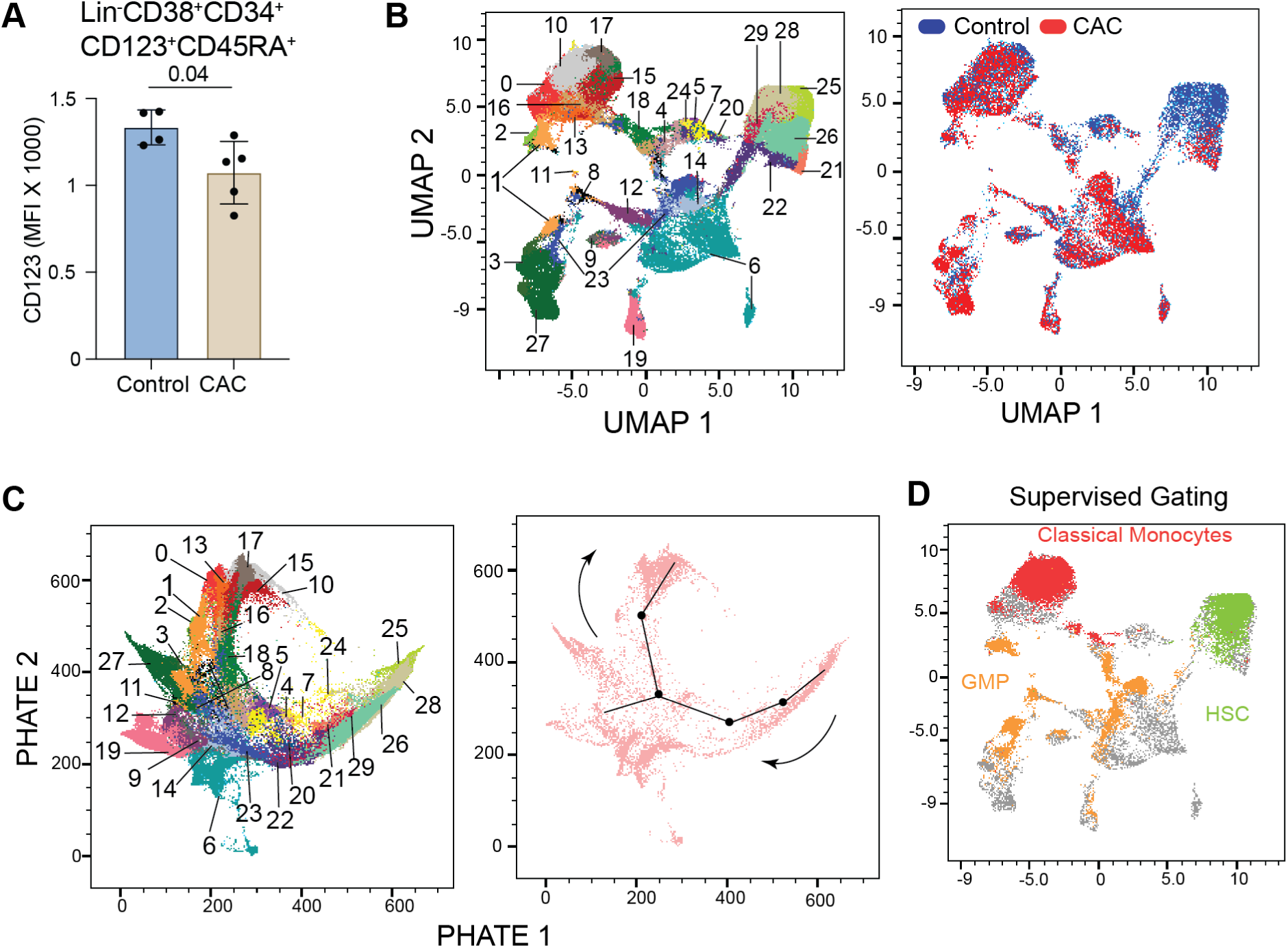
CAC depletes an HSC subset while enhancing a GMP subpopulation. **A**) Expression of CD123 among Lin^-^CD38^+^CD34^+^CD123^+^CD45RA^+^ cells. **B**) UMAP representing identified clusters and contributions of each group. **C**) All clusters were mapped on the PHATE plot. **D**) Monocytes, GMP, and HSC population of supervised analysis were overlapped on the UMAP plot.

**Supp. Fig. 5.**
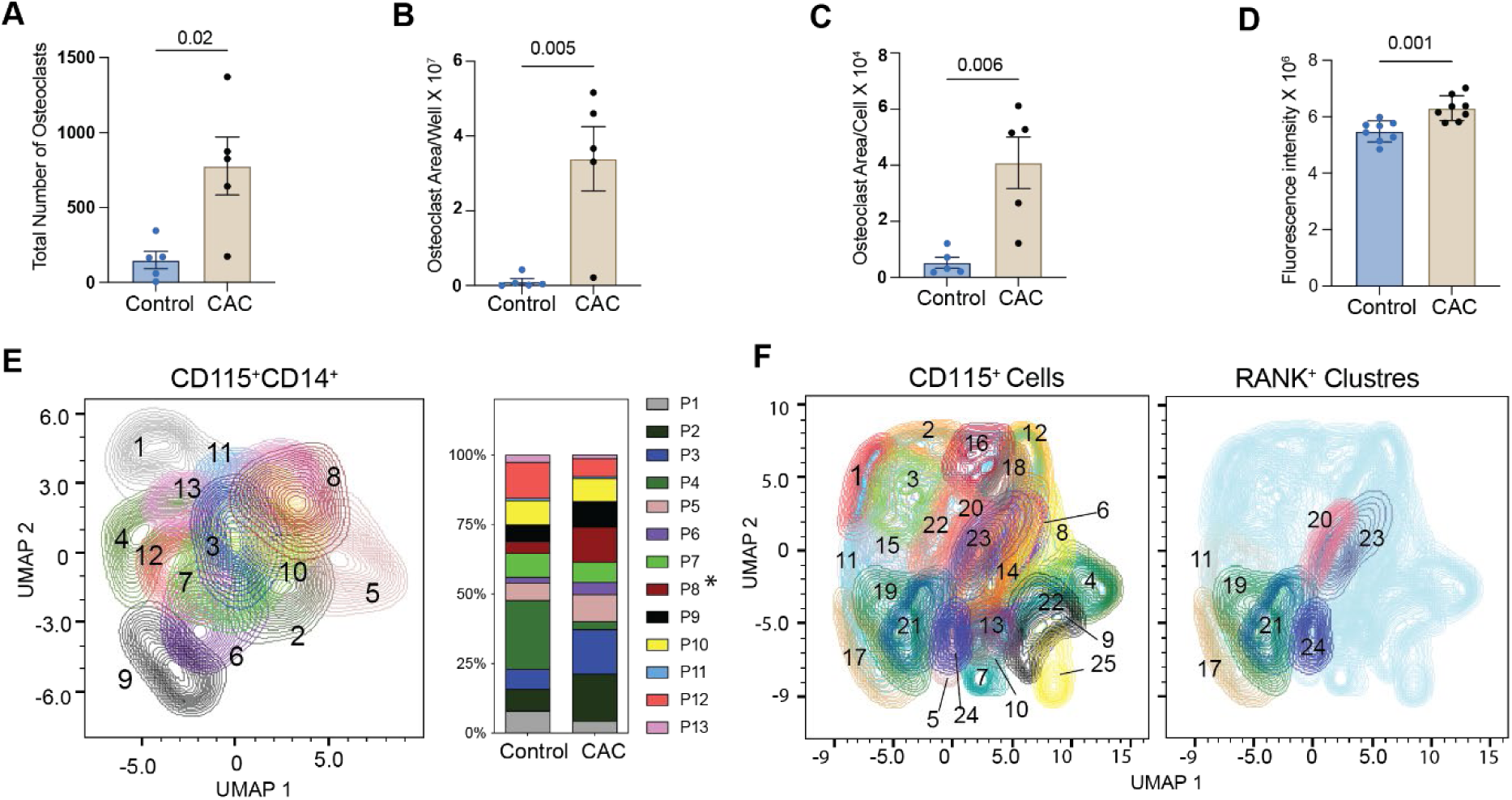
CAC enhances the number and size of osteoclasts. **A)** Total number of osteoclasts, **B**) the osteoclast area per well, and **C)** the osteoclast area per cell identified following *in vitro* osteoclastogenesis. **D**) Quantification of the fluorescence intensity of fluorescein amine-labeled chondroitin in the media of the cells cultured on plates coated with calcium phosphate. **E**) 13 clusters identified among CD115^+^CD114^+^ cells in the panel presented in Fig.2. **F**) Clusters found in CD115^+^ cells within the panel featuring RANK (left), including RANK^+^ clusters (Right).

**Supp. Fig. 6.**
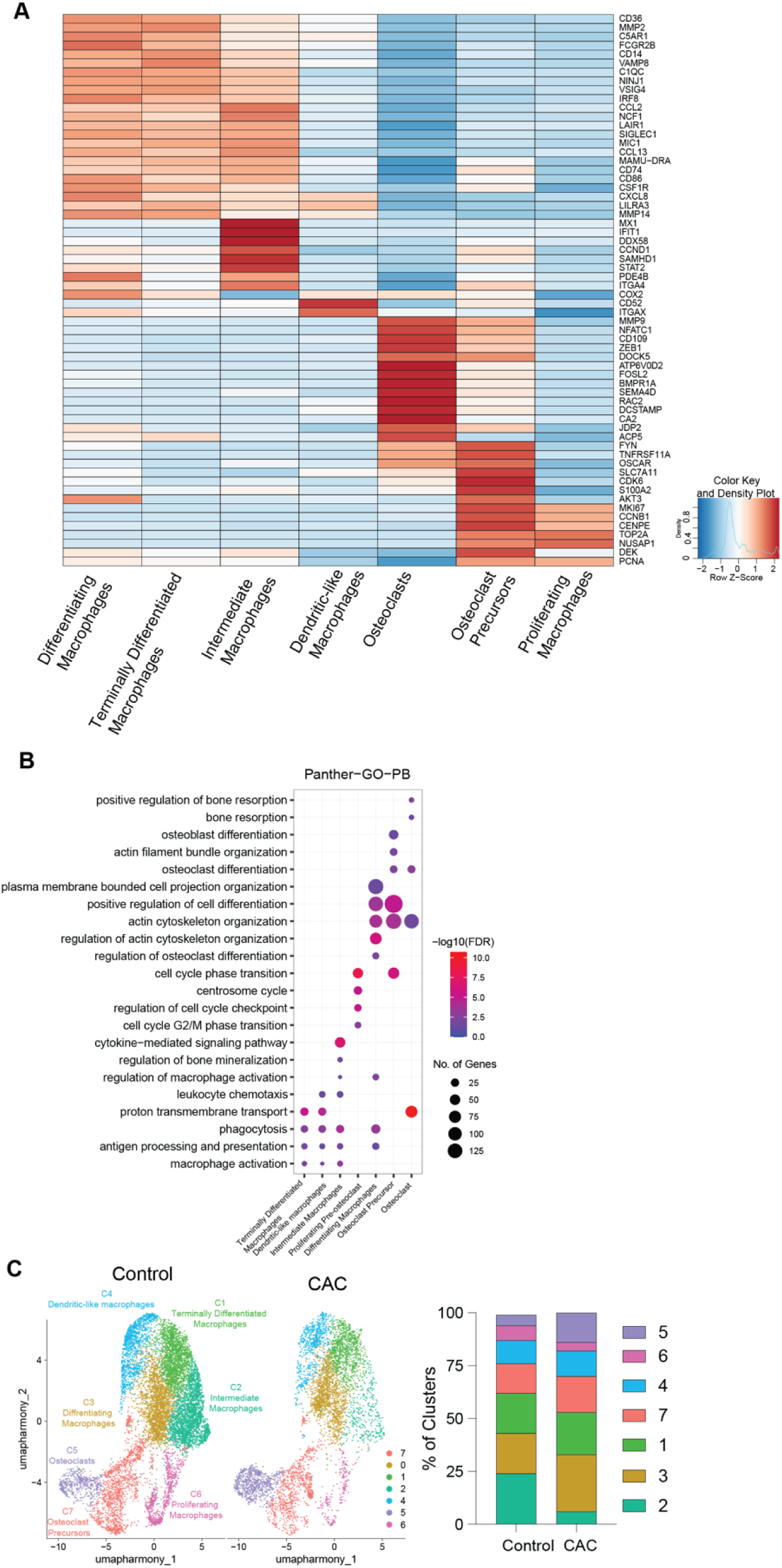
CAC alters the transcriptome profiles of osteoclast precursors. **A**) Heatmap depicting cluster-specific genes. **B**) The cluster genes were enriched for Gene ontology-biological processes using PantherDB. **C**) UMAP depicting the distribution of each cluster across the groups.

**Supp. Fig. 7.**
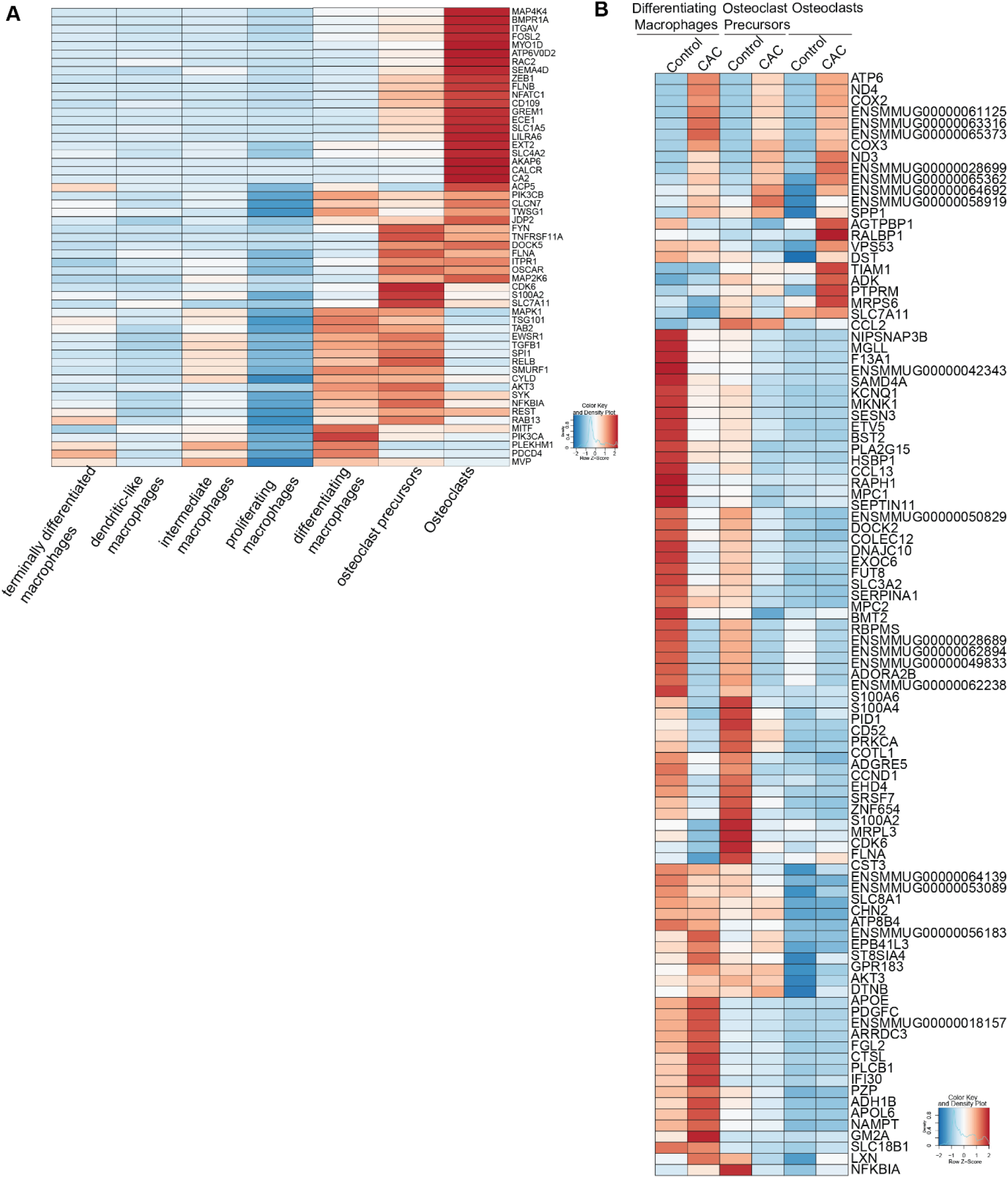
CAC alters the expression of the osteoclast-related genes. **A**) Heatmap illustrating osteoclastogenesis-related genes comparing the expression of genes across clusters. **B**) The list of DEGs across the three osteoclast-related clusters comparing CAC and control.

**Supp. Fig. 8.**
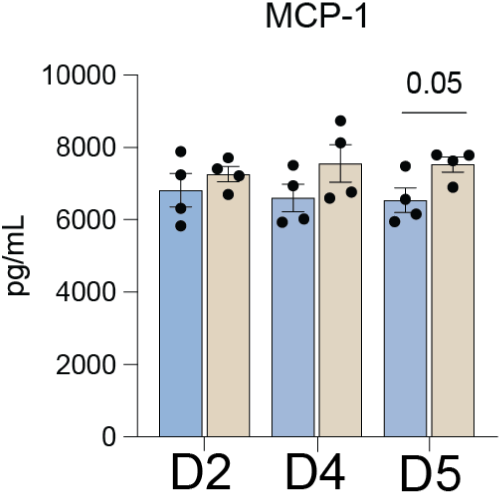
CCL2/MCP-1 was measured in supernatants of the osteoclasts culture using Luminex.

